# Generation and validation of an aryl hydrocarbon receptor knockout human embryonic stem cell line

**DOI:** 10.1101/2025.10.29.685330

**Authors:** Noa Gang, Cuilan Nian, Ekaterina Filatov, Dahai Zhang, Myriam P. Hoyeck, Bailey Laforest, Francis C. Lynn, Jennifer E. Bruin

## Abstract

Glucose homeostasis is tightly controlled by hormones secreted from pancreatic islets. The most abundant cell type in islets is the β-cell, which secretes insulin in response to nutritional stimuli. We previously reported that the adverse metabolic effects of high-dose dioxin exposure in mice are regulated by the aryl hydrocarbon receptor (AHR) specifically in β-cells. Additionally, fetal exposure to low-dose dioxin reduced β-cell area in female mice at birth; however, the role of AHR in β-cell development has not been explored. To characterize the AHR pathway in developing human β-cells, we differentiated human embryonic stem cells (hESCs) into “islet-like” cell clusters (SC-islets) *in vitro* and treated cells with vehicle or dioxin for 24-hours at key stages of differentiation. Dioxin exposure robustly upregulated AHR gene targets (*CYP1A1, AHRR*) at all stages of differentiation but only had modest effects on markers of islet development and maturity. We next generated an AHR knock-out (KO) hESC line and found that basal *CYP1A1* expression was profoundly suppressed in AHR-KO cells compared to parental cells at all stages of differentiation. Key markers of developing and mature pancreatic islets were largely unaffected by AHR deletion; however, *G6PC2* was consistently downregulated in SC-islets from AHR-KO cells compared to parental cells. Interestingly, AHR-KO SC-islets also showed modestly increased insulin secretion relative to the parental line, suggesting a role for AHR in islet development. This novel AHR-KO cell line will allow for deeper investigation into the impact of AHR on development of human islets and other cell lineages.

## 1.0 INTRODUCTION

Type 2 diabetes (T2D) rates are rising globally^1^, yet the underlying factors driving this increase remain unclear. Exposure to persistent organic pollutants (POPs)―including dioxins and dioxin-like chemicals―is consistently associated with increased T2D incidence^2–14^, suggesting that POP exposure is a T2D risk factor. Importantly, there is mounting evidence from experimental model systems showing that POPs detrimentally impact insulin secretion, a critical regulator of glucose homeostasis^15^.

The aryl hydrocarbon receptor (AHR) is a transcription factor originally identified as a receptor for xenobiotics, however, AHR can be activated by both endogenous and exogenous ligands^16^. The most potent AHR ligands are pollutants such as 2,3,7,8-tetrachlorodibenzo-p-dioxin (TCDD or “dioxin”), dioxin-like polychlorinated biphenyls (PCBs), and polycyclic aromatic hydrocarbons (PAHs)^16,17^. In the canonical genomic pathway, ligand-bound AHR translocates to the nucleus where it complexes with aryl hydrocarbon receptor nuclear translocator (ARNT) and binds to xenobiotic response elements (XRE) to induce transcription of target genes, including phase I (e.g., cytochrome P450 1A1 (*CYP1A1*)*, CYP1A2*) and phase II (e.g., glutathione S-transferase A1 (*GSTA1*), NAD(P)H quinone dehydrogenase 1 (*NQO1*)) xenobiotic metabolism enzymes^17–19^. AHR also induces expression of its own repressor protein, aryl hydrocarbon receptor repressor (*AHRR*), which can dimerize with ARNT and subsequently inhibit AHR activity^20^. Importantly, AHR biology is complex and extends far beyond just xenobiotic metabolism^17,21,22^. AHR crosstalks with signaling pathways involved in cell cycle control, lipid metabolism, circadian rhythm, inflammation and immune responses, and adaptation to hypoxia and oxidative stress^22–24^. While most research focuses on the role of AHR in toxicology, there are also important roles for basal AHR activity in cell development and homeostasis^21,22^.

Our lab has shown that 10 nM TCDD exposure robustly activates the AHR pathway—including upregulating *CYP1A1* and *AHRR*—in mouse and human pancreatic endocrine cells (i.e., islets) *in vitro*^25,26^. We also reported that 20 μg/kg TCDD decreased fasting insulin levels and impaired glucose tolerance in mice *in vivo*^27,28^. Importantly, pancreatic β-cell-specific *Ahr* knock-out (KO) mice were protected from the adverse metabolic effects of high-dose TCDD exposure *in vivo*, indicating that AHR activation within β-cells mediates TCDD-induced metabolic dysfunction in adult mice^28^.

In rodent models, global and tissue-specific AHR deletion disrupts the development and function of many organ systems^29–32^, yet the role of AHR in pancreas or islet development has not been reported. Embryogenesis and the early neonatal period are critical windows for β-cell development. β-cell mass is largely established by weaning (i.e., postnatal day (PND) 21) in mice^33–35^ and ∼2 years of age in humans^35,36^. Beyond this developmental window, there is limited capacity for β-cell growth and regeneration^34,37^. Consequently, early life stressors that reduce β-cell mass during infancy are linked to dysregulated metabolic homeostasis in adulthood^38^. Since lipophilic pollutants cross the placenta^39,40^ and are detectable in breast milk^41–47^, offspring exposure begins *in utero*. It is therefore essential to understand how endogenous and exogenous factors influence the formation and maturation of pancreatic β-cells during critical windows of development. Our lab has shown that fetal TCDD exposure caused transient hypoglycemia in female and male mouse pups at PND1 (i.e., birth)^48^. Notably, TCDD-exposed female pups also had decreased β-cell area at PND1, a phenotype that persisted into adulthood^48^. This study suggests that chronic AHR activation *in utero* may impact islet development, but more detailed mechanistic studies are needed.

Human embryonic stem cell (hESC) differentiation provides a powerful *in vitro* model to study fetal development of various cell lineages, including pancreatic islets^49^. Current protocols differentiating hESCs towards the pancreatic β-cell lineage can produce stem cell-derived islets (SC-islets) containing ∼35-70% insulin-producing cells, though their glucose-stimulated insulin secretion capacity remains limited^50,51^. Previous work showed that transient TCDD exposure to undifferentiated hESCs impaired subsequent differentiation towards the pancreatic lineage^52^. Specifically, TCDD reduced the expression of definitive endoderm (*SOX17*) and pancreatic progenitor (*PDX1*) markers^52^, suggesting a possible role of AHR in early pancreatic development. However, this study restricted TCDD exposure to undifferentiated hESCs; the impact of AHR activation at specific differentiation stages remains unknown.

To investigate the role of AHR in pancreatic differentiation, we used a genetically modified hESC reporter line with enhanced green fluorescent protein (eGFP) knocked-in downstream of the insulin promoter (INS-2A-EGFP)^50,51^. We first characterized basal gene expression of key components of the canonical AHR pathway in INS-2A-EGFP cells throughout differentiation toward the β-cell lineage. Next, we exposed INS-2A-EGFP cells to vehicle or 10 nM TCDD for 24-hours at selected differentiation stages to determine (i) whether AHR target genes can be activated at specific stages of pancreatic development, and (ii) the effect of an exogenous AHR ligand on expression of key β-cell lineage markers. Lastly, we generated an AHR-KO INS-2A-EGFP hESC line to investigate the impact of AHR deletion on β-cell differentiation.

## 2.0 METHODS

### 2.1 hESC culture

Undifferentiated INS-2A-EGFP H1 cells^51^ were maintained on Geltrex (Life Technologies, #A1413301) or Matrigel (Corning, #356231) coated plates in mTeSR Plus media (Stem Cell Technologies, #100-0274). Cells were split 1-2 times per week with 6 mL of ReLeSR (Stem Cell Technologies, #100-0484) at approximately 1:10 and plated on 10 cm plates (Corning, #430167). Cells were maintained at 37°C, 21% O_2_, and 5% CO_2_. Parental hESC cells were used in **Figures 1-5**.

**Figure 1.**
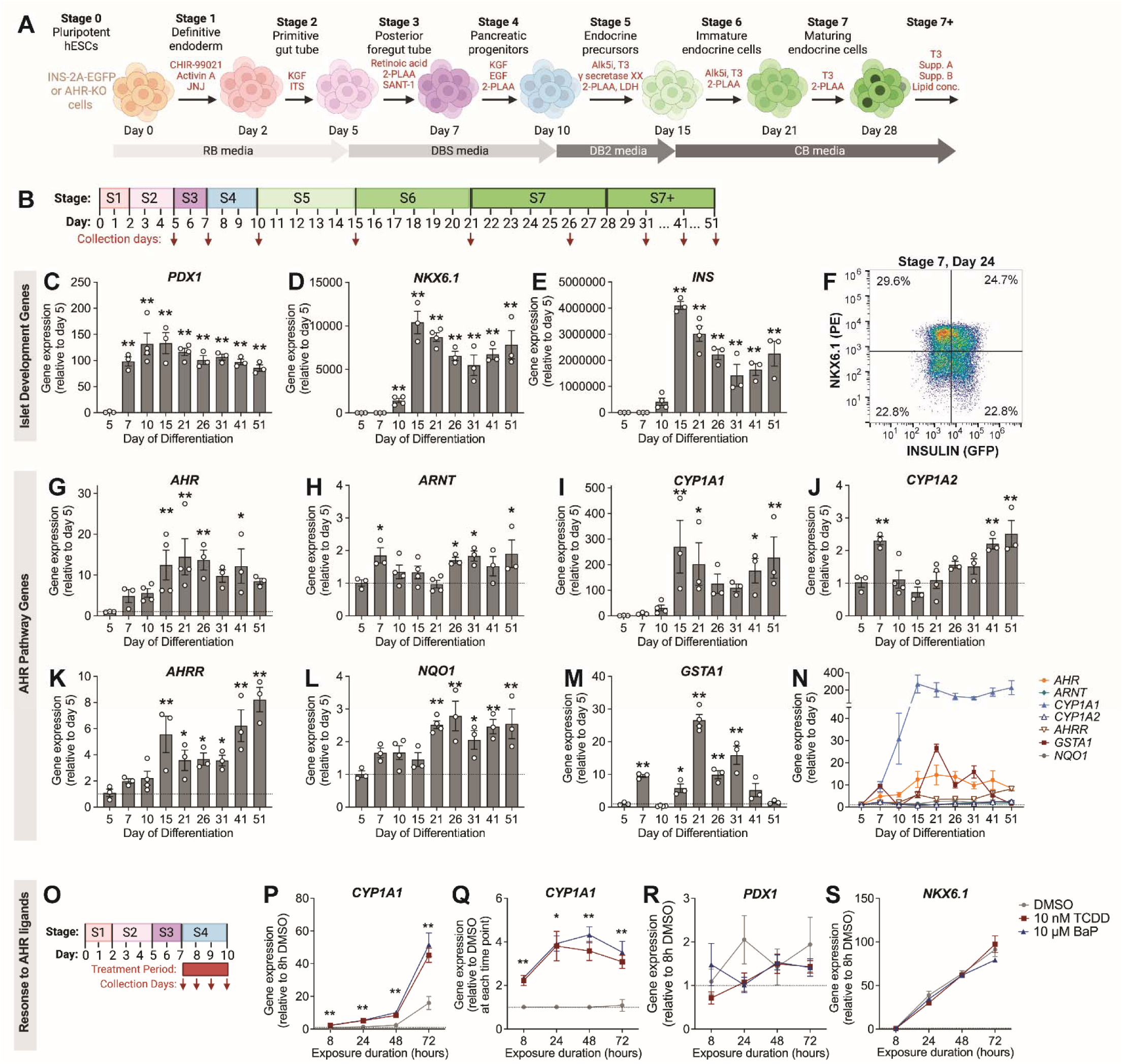
Dynamic expression of key islet development and AHR pathway genes throughout pancreatic differentiation. **(A)** Summary of the 28+ day, 7-Stage differentiation protocol used to generate SC-islets from hESCs. (**B**) Schematic illustrating the days of differentiation where INS-2A-EGFP parental cells were collected for qPCR analysis of basal gene expression. **(C-E)** Gene expression of islet development markers: **(C)** *PDX1,* **(D)** *NKX6.1* and **(E)** *INS*. **(F)** Representative flow cytometry dot plot for Stage 6 cells (Day 24) expressing insulin (GFP) and NKX6.1. **(G-N)** Expression of key AHR pathway genes: **(G,N)** *AHR*, **(H,N)** *ARNT*, **(I,N)** *CYP1A1*, **(J,N)** *CYP1A2,* **(K,N)** *AHRR*, **(L,N)** *NQO1*, and **(M,N)** *GSTA1.* **(C-E,G-N)** Gene expression at all stages of differentiation is normalized to Day 5. Data represent mean +/- SEM and individual data points represent biological replicates (n=3 biological replicates per time point). Asterisks indicate statistically significant differences at each time point compared to Day 5 (*p<0.05, **p<0.01), as determined by one-way ANOVA with Fisher LSD post-hoc test. **(O)** Schematic illustrating the timing of treatment with two AHR ligands (10 nM TCDD, 10 µM BaP) and collection of cells at various timepoints throughout Stage 4. **(P-S)** Gene expression of *CYP1A1, PDX1,* and *NKX6.1* in response to AHR ligands, normalized to DMSO condition at 8 hours **(P,R,S)** or to DMSO at each timepoint **(Q)**. Data represent mean +/- SEM (n=4 technical replicates per time point; n=1 biological replicate). Asterisks indicate statistically significant differences at each time point compared to DMSO (*p<0.05, **p<0.01), as determined by a one-way ANOVA with Dunnett post-hoc test. (**A,B,O**) Schematics were made with BioRender.com.

### 2.2 Guide and modified AHR donor vector construction

The parental INS-2A-EGFP H1 line^51^ was used to generate the AHR-KO clones. In brief, the CRISPR/Cas9 vector was based on px458 (Addgene; plasmid 48138), but with the exchange of a Cdh to a full-length CAGGS promoter to maximize hESC expression (pCCC)^53^. The gRNA (CCT ACG CCA GTC GCA AGC GG) was designed using the algorithm reported by Doench *et al.*^54^ and cloned into the BbsI sites of pCCC to generate pCCC-LL1194 as previously described^55^.

AHR 5’ and 3’ homology arms (both 800 bp) were amplified from H1 genomic DNA using a proofreading polymerase (Hercules Fusion) and cloned into KpnI/NheI and AscI/NotI sites, respectively. A Geneblock was ordered that included an HA-epitope tag for knock-in identification^56^ and FKBP12^F36V^-mScarlet fusion into the first exon of *AHR* for potential inducible degradation in the future^57^ (**Supplemental Table 1**^58^). The Geneblock sequence was cloned into the NheI/EcoRI sites^57^. However, we did not use the dTAG molecular control system in this study. The Floxed PGK-Puro cassette used for clone selection contained a polyadenylation termination sequence that facilitated premature (first exon) termination of *AHR* transcription and was sufficient to render any AHR protein translated incomplete and dysfunctional, essentially “knocking out” the AHR (**Figure 3A**). The constructs were fully Sanger sequenced in-house prior to transfection.

### 2.3 Generation of the AHR-KO cell line

INS-2A-EGFP cells were transfected at 30-40% confluency and fed with mTesR Plus media (Stem Cell Technologies, #05825) at least 1 hour prior to transfection. Cells were washed with PBS without calcium and magnesium (-Ca^+2^/Mg^+2^; Cytiva, #SH30256.01) before lifting with Accutase (Life Technologies, #07922) for 6 minutes at 37 °C to generate a single cell suspension. Following detachment, cells were centrifuged at 200 x g for 5 minutes before being washed with PBS (+ Ca^+2^/Mg^+2^; Gibco, #14040-133). INS-2A-EGFP cells (5x10^6^ cells) were resuspended in 700 μL of room temperature PBS (+ Ca^+2^/Mg^+2^) and transferred to a 0.4 cm gap cuvette containing 40 µg of pAHR-FKBP12 ^F36V^-mScarlet-PGKPuro donor plasmid and 15 µg of guide plasmid (pCCC-LL1194). The cuvette containing cells and plasmids was electroporated using Bio-Rad Gene Pulser II system (250 V, 500 µF, infinite Ω, time constant = 9.3 msec). After electroporation, cells were resuspended into StemFlex (Gibco, #A33493.01) with 10 µM Y-27632 dihydrochloride (Selleck Chemicals, #S6390) and plated onto a 6 cm Laminin-521 (diluted to 12% in PBS; BioLamina, #LN521-05) coated plate. Media was replaced daily, and cells were allowed to recover for 3 days before selecting with 0.375 μg/mL puromycin (Sigma-Aldrich, #P8833). Surviving colonies were picked into a 96-well Laminin-coated plate within 7 days of electroporation by manual scraping and pipetting the colony off the plate and into a 96 well Geltrex-coated plate with 100 μL StemFlex medium (Gibco, #A3349401). Once clones were close to confluent, the plate was split into 3, 96-well Geltrex-coated plates with 100 μL mTeSR Plus per well. One plate was genotyped and 2 were frozen down to expand the correctly targeted clones.

Genomic DNA was extracted using QuickExtract (Mandel, #QE09050). Primers were designed to match the AHR 5’ and 3’ homology arms (**Supplemental Table 2**^58^). Extracted genomic DNA was run on a 1% agarose (Fisher Scientific, #BP160-500) gel in buffer TAE made in-house (40 mM Tris base from Fisher Bioreagents, #BP152-5; 0.1% acetic acid from Fisher Scientific, #351272-212; and 1 mM EDTA from Fisher Scientific, #15575020) and visualized using Redsafe (FroggaBio, #21141; 1:20,000 dilution). Genotyped clones containing both 5’ and 3’ AHR arms and an absence of wild type (WT) AHR were expanded, frozen at passage numbers 37-40, and Sanger sequenced in-house. Three clones identified to possess the correct genotype were differentiated to Day 9 (see Section 2.4) to verify the absence of AHR protein by Western blot. The hPSC Genetic Analysis Kit (Stem Cell Technologies, #07550) was used as per the manufacturers recommendations to assess for the presence of common recurrent karyotypic abnormalities reported in hESC lines. AHR-KO cells were used in **Figures 3-5**.

### 2.4 In vitro differentiation of hESCs into SC-islets

Parental and AhR-KO hESCs were differentiated according to a 7-Stage suspension protocol adapted from Santini-González *et al*^59^ (summarized in **Figure 1A** and **Supplemental Table 3**^58^). Cells were maintained at 37°C and 21% O_2_. To initiate differentiation, hESCs were lifted with 3 mL of Accutase for 6 minutes at 37°C to generate a single cell suspension. Following detachment, cells were diluted 1:3 in PBS (+ Ca^+2^/Mg^+2^) and centrifuged at 200 x g for 5 minutes before being resuspended at 1 x 10^6^ cells/mL in mTeSR Plus supplemented with 10 μM Y-27632 dihydrochloride. Cells were seeded into an uncoated 6-well plate (Fisher Scientific, #657185) at 5.5 mL/well (5.5 x 10^6^ cells/well) and placed on the orbital shaker (95 or 105 RPM) for at least 24 hours to promote cluster formation. Prior to initiating Stage 1 of differentiation, cells were washed twice with 5.5 mL of PBS (+ Ca^+2^/Mg^+2^) for 10 minutes on the orbital shaker.

The specific additives used throughout the 7-Stage differentiation protocol are specified in **Supplemental Tables 3 and 4**^58^. The base media formulations are as follows (all with a pH of 7.4): **(a) “RB Media”**: RPMI 1640 media (Fisher Scientific, #SH3002701) supplemented with 0.5X penicillin/streptomycin (HyClone, #SV30010) and 1X glutaMAX-I (Gibco, #35050-061); **(b) “DBS media”**: DMEM high glucose media (Fisher Scientific, #SH30081.01) supplemented with 0.5X penicillin/streptomycin, 1X glutaMAX, 1X non-essential amino acids (NEAA; Gibco, #11140-050 or Stem Cell Technologies, #07600), and 1 mM sodium pyruvate (Gibco, #11360070); **(c) “DB2 media”**: DBS media plus 1% fatty-acid free bovine serum albumin (BSA; Proliant Biologicals, #68700 or Sigma-Aldrich, #A7030) and 1X ITS; **(d) “CB media”**: CMRL 1066 media (United States Biological, #C5900-01) supplemented with 1% BSA, 1X GlutaMAX, 0.5% Pen/strep, 1 mM sodium pyruvate, 1X ITS, 1X NEAA, 10 μM Zinc Sulfate (Sigma-Aldrich, #Z0251-100G), 1X HEPES (Sigma-Aldrich, #H3375-500G), 1 mM N-acetyl-L-cysteine (NAC; Sigma-Aldrich, #A9165-25G), 10 μg/mL Heparin (Sigma-Aldrich, #H3149-50KU), and 0.05 mM 2-mercaptoethanol (2-ME; Sigma Aldrich, #M7522-250ML). with a pH of 7.4.

Cell cluster diameter was assessed on Day 30 for parental cells and all AHR-KO clones (n = 100 technical replicates) using ImageJ Software.

### 2.5 Chemical treatment

Schematics are provided alongside each figure to clearly illustrate the timing for chemical treatment and cell collection. In the first experiment, ∼100 INS-2A-EGFP clusters were exposed to DMSO (vehicle control; Sigma-Aldrich, #D2650), 10 nM TCDD (Accustandard, #D-404S-DMSO-10x), or 10 µM benzo(a)pyrene (BaP; Sigma-Aldrich, #B1760-100MG) for up to 72 hours starting on Day 7 of differentiation. A subset of cells were collected after 8, 24, 48, and 72 hours of treatment for analysis by qPCR (**Figure 1O**; n=4 technical replicates per time point, n=1 biological replicate). Next, ∼40 INS-2A-EGFP clusters were exposed to DMSO or different doses of TCDD (1, 10, 100 nM) for 24 hours starting on Day 14 of differentiation (**Supplemental Figure 1**^58^; n=4 technical replicates per dose, n=1 biological replicate). For these experiments, clusters were treated on the orbital shaker (115 RPM) in 24-well plates (Thermo Fisher Scientific, #144530) containing 1 mL of differentiation media.

To examine AHR activation at specific stages of differentiation, INS-2A-EGFP cells (n=3 biological replicates, i.e., separate differentiations) were removed from the orbital shaker and exposed to DMSO or 10 nM TCDD for 24 hours starting on the following days of differentiation: Day 4 (end of Stage 2; **Figure 2A**), Day 6 (end of Stage 3; **Figure 2B**), Day 9 (end of Stage 4; **Figure 2C**), Day 14 (end of Stage 5; **Figure 2D**), Day 20 (end of Stage 6; **Figure 2E**), Day 25 (mid Stage 7; **Figure 2F**), and Day 40 (mid Stage 7^+^; **Figure 2G**). AHR-KO cell lines were treated similarly but starting on Day 9 only (Stage 4; **Figure 3D**). For these experiments, ∼200-400 clusters were treated in 6-well plates (Cellstar, #657185) containing 5 mL of differentiation media.

**Figure 2.**
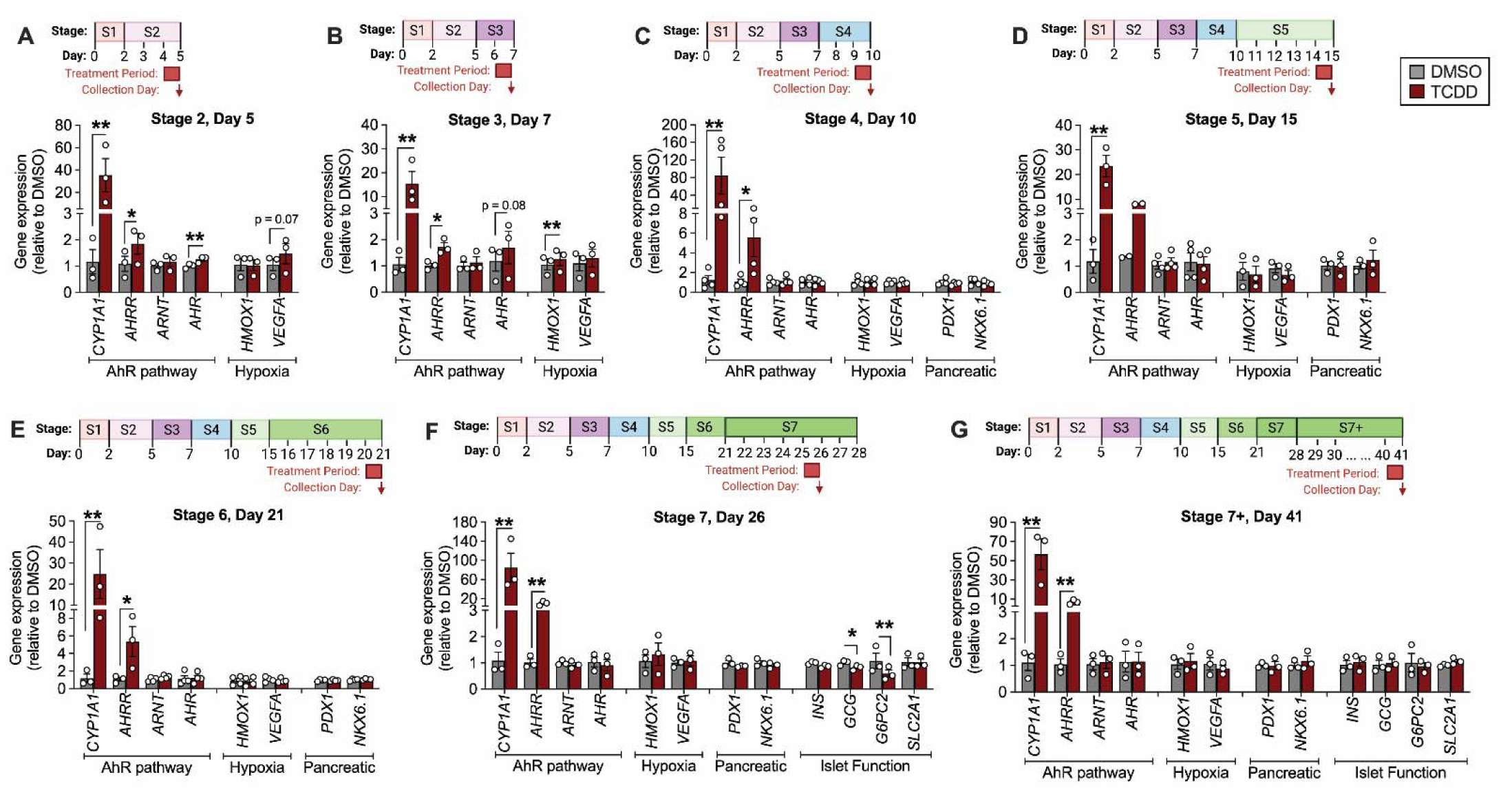
TCDD causes upregulation of *CYP1A1* and *AHRR* expression at all stages of pancreas differentiation. INS-2A-EGFP parental hESCs were exposed to either vehicle (DMSO) or TCDD (10 nM) for 24 hours at the indicated days throughout the pancreatic differentiation protocol. Schematics are provided to clearly illustrate the timing of treatment and collection of cells. Gene expression is normalized to DMSO at each timepoint: **(A)** Day 5 (Stage 2), **(B)** Day 7 (Stage 3), **(C)** Day 10 (Stage 4), **(D)** Day 15 (Stage 5), **(E)** Day 21 (Stage 6), **(F)** Day 26 (Stage 7), and **(G)** Day 41 (Stage 7+). Data represent mean +/- SEM and individual data points represent biological replicates (n=3 biological replicates per time point). Asterisks indicate statistically significant differences between vehicle vs TCDD-exposed cells (*p<0.05, **p<0.01), as determined by a ratio paired t-test. Schematics were made with BioRender.com.

Following each treatment period, cells were rinsed 3 times with PBS (- Ca^+2^/Mg^+2^) (Cytiva, #SH30264.01), resuspended in buffer RLT (Qiagen) + 10% 2-ME, and stored at -80°C until real time quantitative polymerase chain reaction (RT-qPCR) analysis.

### 2.6 RT-qPCR

mRNA was isolated using the RNeasy Micro kit (Qiagen, #74004) as per the manufacturer’s instructions. DNase treatment was performed prior to cDNA synthesis with the iScript^TM^ gDNA Clear cDNA Synthesis Kit (Biorad, #1725035). RT-qPCR was performed using SsoAdvanced Universal SYBR Green Supermix (Biorad, #1725271) and run on a Vii7 instrument (Applied Biosystems) or a CFX384 instrument (Biorad). *PPIA* was used as the reference gene. Primer sequences are listed in **Supplementary Table 5**^58^. Data were analyzed using the 2^-ΔΔCT^ relative quantitation method.

### 2.7 Western blot

Unexposed parental and AHR-KO cell lines were collected for Western blots on Day 10 (**Figure 3C**). Cells were rinsed in PBS (+ Ca^+2^/Mg^+2^) and resuspended in lysis buffer (2% SDS and 10% glycerol). Protein was denatured by boiling at 95°C for 10 minutes before lysis by sonication (S-4000 with cuphorn; Misonix) for 2 minutes. Cells were then centrifuged at 200 x g for 5 minutes at 20°C and supernatant was collected. Lysates were subjected to standard SDS-PAGE (30% acrylamide, 10% SDS) followed by wet transfer blotting onto a nitrocellulose membrane (Biorad, #1620177). Blots were then blocked with 5% milk powder in Tris-buffered saline with 0.1% Tween (Thermo Fisher Scientific, #BP337-500) and probed with horseradish peroxidase conjugated mouse anti-human AHR (Santa Cruz Biotechnology Incorporated, sc-133088, RRID:AB_2273721; 1:100), anti-HA epitope (Abcam, #AB9110, RRID:AB_307019; 1:1000) or mouse anti-GAPDH (Sigma-Aldrich, #G8795-100UL, RRID:AB_1078991; 1:125,000) overnight at 4°C in blocking buffer. Membranes were then washed and probed with horseradish peroxidase conjugated goat anti-mouse secondary antibodies (Jackson ImmunoResearch, #115-035-174, RRID:AB_2338512; 1:1000) for 1 hour and visualized with Immobilon Forte HRP Substrate (Sigma-Aldrich, #WBLUF0500).

**Figure 3.**
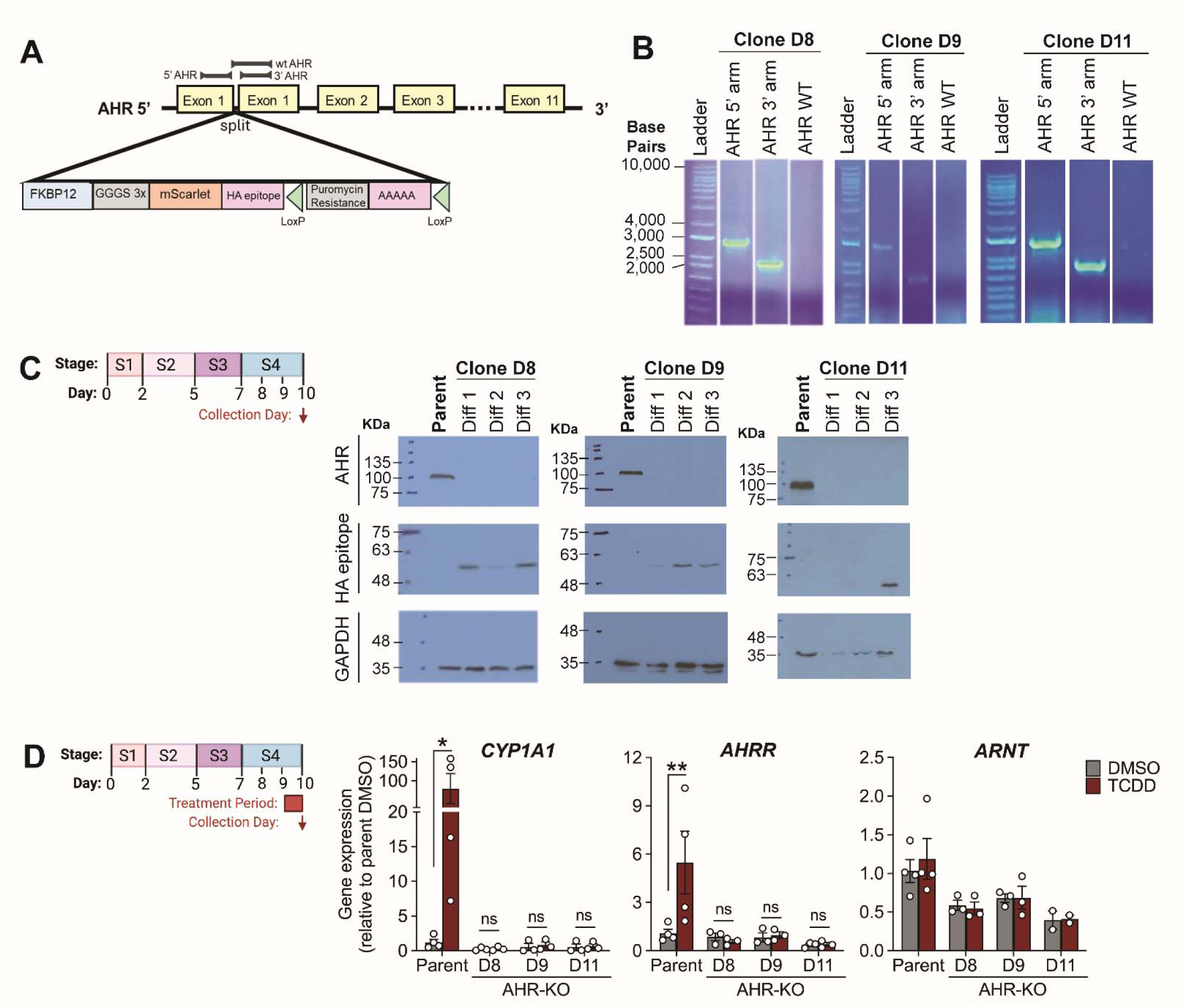
Generation and validation of AHR-KO cell line. **(A)** AHR was modified at a split in the first exon, with an early poly-adenyl tail insert, truncating transcription. Primer sites for *AHR* 5’ and 3’ homology arms and wildtype (WT) *AHR* are shown. **(B)** AHR-KO clones D8, D9, and D11 were assessed for the presence of 5’ and 3’ *AHR* arms and the WT *AHR* sequence. All 3 clones were validated to be positive for 5’ and 3’ *AHR* homology arms and homozygous for the absence of WT *AHR* genomic DNA. **(C)** Untreated INS-2A-EGFP parent cells and AHR-KO clones were collected for Western blot analysis on Day 10 of differentiation. Protein expression was measured for AHR, the HA epitope, and housekeeper GAPDH in the INS-2A-EGFP parent line (n=1 biological replicate) and AHR-KO clones D8, D9, and D11 (n=3 biological replicates per clone; i.e. “Diffs 1, 2, 3”). **(D)** INS-2A-EGFP parental cells (n=4 biological replicates) and AHR-KO clones (n=3 biological replicates per clone) were differentiated to Day 9 and then exposed to either vehicle (DMSO) or TCDD (10 nM) for 24 hours. Gene expression of *CYP1A1, AHRR,* and *ARNT* was measured and normalized to the parental line DMSO control. Data represent mean +/- SEM and individual data points represent biological replicates. Asterisks indicate statistically significant differences (*p<0.05, **p<0.01), as determined by two-way repeated measures ANOVA with Sidak post-hoc test. Schematics were made with BioRender.com.

### 2.8 Flow cytometry

Unexposed parental (Day 40-52) and AHR-KO SC-islets (Day 43-49) were collected for flow cytometry analysis (n=3-5 biological replicates per genotype). SC-islets were transferred to 15 mL conical tubes and cells were pelleted at 1300 rpm for 5 minutes. The cell pellet was washed once with pre-warmed PBS (+ Ca^+2^/Mg^+2^), dispersed in 400 μL Accutase (Corning, #25-058-CI) and the reaction was quenched with 600 μL of CB media. Cells were pelleted at 1300 rpm for 5 minutes, washed with PBS (+ Ca^+2^/Mg^+2^), and again pelleted to remove the supernatant. Cells were then resuspended in 200 μL of PBS (+ Ca^+2^/Mg^+2^) and transferred to a 96-well flat bottom plate. GFP fluorescence was measured using the fluorescence 1 channel on a BD Accuri C6 flow cytometer using BD Biosciences Accuri C6 software (BD Biosciences, Mississauga, ON). The %GFP^+^ cells was calculated as %GFP^+^ cells / total cells counted, with 20,000 events per sample. Refer to **Supplemental Figure 1**^58^ for gating strategy.

To quantify NKX6.1 and GFP (i.e., insulin) protein, INS-2A-EGFP SC-islets were collected at the end of stage 6 (Day 24). SC-islets were transferred to 1.5 mL tubes, washed once with pre-warmed PBS (+ Ca^2+^/Mg^2+^) and dispersed in 500 μL Accutase (Sigma-Aldrich, #A6964) for 10 minutes at 37°C, with firm tapping every 2 minutes. The reaction was quenched with 1 mL of cold flow buffer (PBS (-Ca^+2^/Mg^+2^), 1% BSA, 2 mM EDTA) and cells were filtered into 5 mL flow tubes with a 35 μm strainer cap (Corning, #352235). Cells were pelleted at 200 x g for 5 minutes, washed twice with 1 mL of cold PBS (-Ca^+2^/Mg^+2^), resuspended in 100 μL of LIVE/DEAD Fixable Far Red Dead Cell Stain Solution (1:1000 dilution in PBS (-Ca^+2^/Mg^+2^); Invitrogen, #L34974), and incubated at 4°C for 30 minutes in the dark. Cells were then washed once with 1 mL of PBS (-Ca^+2^/Mg^+2^), once with 1 mL flow buffer, and fixed in 250 μL of Cytofix/Cytoperm Fixation and Permeabilization Solution (BD Biosciences, #554722) at 4°C for 20 minutes in the dark. Cells were then washed twice with 1 mL of Perm/Wash Buffer (BD Biosciences, #554723), resuspended in 100 μL of Perm/Wash Buffer containing PE-anti-NKX6.1 (BD Biosciences, #563023, RRID:AB_2716792; 1:20 dilution), and incubated at 4°C for 30 minutes in the dark. Cells were then washed twice with 1 mL of Perm/Wash Buffer, resuspended in 200 μL of flow buffer, and transferred to 1.5 mL tubes. Data were acquired on an Attune NxT Flow Cytometer (Invitrogen) with a minimum of 10,000 events collected per sample. Dot plots were created with FlowJo v10. Refer to **Supplemental Figure 2**^58^ for gating strategy.

### 2.9 Perifusion analysis

Unexposed parental and AHR-KO SC-islets (Day 26-55; see **Figure 5G** for exact age of each biological replicate) were collected to measure dynamic insulin secretion by perifusion analysis (n=5-6 biological replicates per genotype; Biorep Technologies, Miami Lakes, FL). SC-islets from all three AHR-KO clones were included in the perifusion analysis (n=4 from clone D8, n=1 from clone D9, n=1 from clone D11; see **Figure 5I**). For each biological replicate, 70 SC-islet clusters were hand-picked and loaded into Perspex microcolumns sandwiched between two layers of acrylamide-based microbeads (Biorep Technologies, #PERI-BEADS-20). Clusters were perifused for 40 minutes with 2.8 mM low glucose (LG) KRBB (115 mM NaCl, 5 mM KCl, 24 mM NaHCO_3_, 2.5 mM CaCl_2_, 1 mM MgCl_2_, 10 mM HEPES, 0.1 % (wt/vol.) BSA, pH 7.4) at a rate of 40 μL/min to equilibrate the clusters and the effluent was discarded. Clusters were then perifused with the following protocol: 25 minutes with 2.8 mM LG-KRBB at a flow rate of 40 μL/min, 12.5 minutes with 16.7 mM high glucose (HG) KRBB at a flow rate of 80 μL/min, 15 minutes with HG-KRBB at a flow rate of 40 μL/min, 20 minutes with LG-KRBB at a flow rate of 40 μL/min, 15 minutes with 30 mM KCl-KRBB at a flowrate of 80 μL/min, 20 minutes with KCl-KRBB at a flowrate of 40 μL/min, and 25 minutes with LG-KRBB at a flowrate of 40 μL/min. Throughout the protocol, the SC-islets and perifusion solutions were maintained at 37°C using the built-in temperature-controlled chamber, while the collection plate was cooled to 4°C with the tray cooling system. Samples were stored at -80°C until analysis. Insulin concentrations were measured by radioimmunoassay (Millipore, KI-14K, RRID:AB_2801577) and replicates across experiments were pooled. Area under the curve (AUC) was calculated as follows: LG AUC from 5-25 minutes; HG AUC from 27.5-67.5 minutes, and KCl AUC from 75-107.5 minutes.

### 2.10 Statistics

All statistics were performed using GraphPad Prism 10.4.1 (GraphPad Software Inc., La Jolla, CA). Specific statistical tests are defined in the figure legends. For all analyses, p<0.05 was considered statistically significant and any potential outliers were tested for significance using a Grubbs’ test with α=0.05. Non-parametric statistics were used in cases where the data failed normality or equal variance tests. Data are presented as mean ± SEM.

## 3.0 RESULTS

### 3.1 The AHR pathway is basally induced during pancreatic differentiation

We used the parental INS-2A-EGFP cell line to first assess basal expression of AHR pathway genes throughout pancreatic differentiation, alongside key markers of islet development (**Figure 1A,B**). As expected, *PDX1* was robustly induced on Day 7 of differentiation and then subsequently maintained throughout the differentiation (**Figure 1C**). *NKX6.1* was induced starting at Day 10 and then further increased at Day 15 (**Figure 1D**). *INS* was profoundly induced from Day 15 onward (**Figure 1E**). A representative flow cytometry plot is shown to illustrate the efficiency of a typical differentiation, with 47.5% of live cells expressing GFP/insulin and 24.7% co-expressing both GFP/insulin and NKX6.1 at Day 24 of differentiation (end of Stage 6; **Figure 1F**).

Interestingly, markers of the AHR pathway increased shortly after pancreatic endocrine cell induction (**Figure 1G-N**). Basal *AHR* expression was significantly increased starting at Day 15 relative to Day 5 (**Figure 1G**), coinciding with the robust induction of downstream AHR targets *CYP1A1* (mean induction ranging from 110-fold to 270-fold between Days 15-51; **Figure 1I**) and *AHRR* (mean induction ranging from 3.6-fold to 8.2-fold between Days 15-51; **Figure 1K**). *NQO1* had a similar but delayed pattern of expression, increasing by Day 21 (2.6-fold relative to Day 5; **Figure 1L**). Other downstream AHR targets, *CYP1A2* and *GSTA1,* showed biphasic induction on Day 7 (end of Stage 3) and then again starting at either Day 41 for *CYP1A2* (Stage 7+; **Figure 1J**) or Day 15 for *GSTA1* (end of Stage 5; **Figure 1M**). *ARNT,* an important cofactor for AHR, remained relatively stable throughout differentiation (**Figure 1H**). Among all genes examined, *CYP1A1* was by far the most strongly upregulated AHR target gene during pancreatic differentiation (**Figure 1I,N**).

### 3.2 Exogenous AhR ligands can further activate the AHR pathway in pancreatic progenitor cells

To determine if the AhR pathway can be further activated beyond the basal induction seen in Stage 4-5 of pancreatic differentiation, we exposed Stage 4 parental cells to two different exogenous AHR ligands—TCDD and BaP—for up to 72 hours (**Figure 1O**). TCDD was tested at 1000x lower concentration than BaP (10 nM TCDD versus 10 µM BaP), given the known differences in ligand potency^60^. This dose of TCDD was previously shown to upregulate both *CYP1A1* gene expression and CYP1A1 enzyme activity in primary mouse and human islets without causing cytotoxicity^25^. It is important to note that 10 nM TCDD does not maximally activate AHR; we demonstrated that 100 nM TCDD can further amplify *CYP1A1* induction beyond 10 nM TCDD (**Supplemental Figure 3**^58^).

*CYP1A1* gene expression increased basally (i.e., in DMSO conditions) throughout Stage 4 and was further upregulated by both ligands at all timepoints (8, 24, 48, 72 hours; **Figure 1P**). The magnitude of *CYP1A1* induction relative to DMSO was similar at all timepoints for both ligands, ranging from 2.2-fold to 3.6-fold for TCDD and 2.3-fold to 4.3-fold for BaP (**Figure 1Q**). The expression of key pancreatic progenitor cell transcription factors, *PDX1* and *NKX6.1,* was not affected by TCDD or BaP exposure at any timepoint throughout Stage 4 (**Figure 1R,S**).

### 3.3 TCDD activates the AHR pathway at all stages of pancreatic differentiation

To determine if the AHR pathway can be activated by an exogenous ligand throughout pancreatic endocrine cell development, parental INS-2A-EGFP cells were exposed to DMSO or 10 nM TCDD for 24 hours at selected stages of pancreas differentiation (see schematics in **Figure 2** for specific timing). While *AHR, CYP1A1* and *AHRR* were not basally induced until Day 15 (**Figure 1G,I,K**), TCDD robustly upregulated *CYP1A1* and *AHRR* at all stages of pancreatic differentiation examined (**Figure 2A-G, Supplemental Figure 4A,B**^58^). *CYP1A1* was induced up to 165-fold (peak in Stage 4) and *AHRR* up to 14.8-fold (peak in Stage 7) following TCDD exposure compared to vehicle (**Figure 2A-G, Supplemental Figure 4A,B**^58^). TCDD modestly increased expression of *AHR* in Stage 2 (mean induction of 1.3-fold; **Figure 2A**) but did not impact *AHR* expression at other stage of differentiation (**Figure 2B-G, Supplemental Figure 4C**^58^). Given that ARNT is a binding partner for both AHR and HIF1α, we measured *ARNT* and HIF1α targets, *HMOX1* or *VEGFA*; TCDD did not interfere with expression of any of these targets throughout pancreatic differentiation (**Figure 2A-G, Supplemental Figure 4D-F**^58^).

We next measured the impact of 24-hour TCDD exposure on expression of genes related to β-cell development and identity. TCDD did not impact *PDX1* or *NKX6.1* expression at any stage of differentiation (**Figure 2A-G, Supplemental Figure 4G,H**^58^). TCDD modestly but significantly decreased expression of *GCG* and *G6PC2* in Stage 7 (**Figure 2F, Supplemental Figure 4J,K**^58^), but these effects were not consistently seen throughout Stage 7+ of differentiation (**Figures 2F, 2G, Supplemental Figure 4J,K**^58^). *INS* and *SLC2A1* were unaffected by TCDD exposure in Stages 7 or 7+ (**Figure 2F,G; Supplemental Figure 4I,L**^58^).

### 3.4 Generation and validation of the AHR-KO cell line

Given that the AHR pathway was activated during β-cell differentiation both basally (**Figure 1G-N**) and following exposure to an exogenous ligand (**Figure 1P,Q and Figure 2**), we next generated an AHR-KO hESC line to investigate the role of AHR during pancreatic endocrine cell development. We built the AHR-KO construct (**Figure 3A**) as described in the methods. Briefly, the FKBP12^F36V^ degron sequence was inserted into the first AHR exon. Immediately downstream of FKBP12^F36V^, a 12 amino acid Gly-Ser-Gly repeat (GGGS x3) linker connected a mScarlet reporter protein sequence fused to a HA-epitope sequence. The downstream premature polyadenylation signal sequence truncated AHR transcription early within the first exon resulting in non-functional AHR protein.

The parental INS-2A-EGFP cell line was transfected with the AHR-KO construct and cells were selected in puromycin. Clones were expanded and genomic DNA was collected and amplified by qPCR across AHR 5’ and 3’ homology arms, and the WT AHR sequence. Of the clones generated, three (clones D8, D9, and D11) showed a clear PCR product across both 5’ and 3’ AHR homology arms and an absence of WT AHR genomic DNA (**Figure 3B**). All three clones were Sanger sequenced and did not show any off-target mutations (data not shown). Stem Cell Technologies Genetic Analysis Kit was used to look for common chromosomal mutations. Clones D8 and D11 both showed no abnormal chromosomal mutations; clone D9 had a deletion in a minimally important region of Chromosome 17q (**Supplemental Figure 5**^58^).

We next measured expression of AHR protein by Western blot in each of the three AHR-KO clones (D8, D9, D11) and the parental INS-2A-EGFP line at the end of Stage 4 (**Figure 3C**). As expected, the parental cells showed robust expression of AHR protein and no detectable levels of the HA epitope (**Figure 3C**). In contrast, the three AHR-KO clones all had no detectable AHR protein and variable expression of the HA epitope protein (**Figure 3C**). These results strongly suggest successful knockout of the AHR in each of the three clones.

As final validation of the AHR-KO, we treated the parental cells and AHR-KO clones with DMSO or 10 nM TCDD for 24 hours starting on Day 9 of differentiation to determine whether TCDD exposure would induce downstream AHR targets (**Figure 3D**). As expected, in parental cells, TCDD exposure significantly induced *CYP1A1* (induction ranging from 7.2-fold to 159-fold) and *AHRR* (induction ranging from 1.9-fold to 10.1-fold) relative to DMSO (**Figure 3D**). Importantly, there was no upregulation of *CYP1A1* or *AHRR* by TCDD in any AHR-KO clones (**Figure 3D)**. TCDD did not impact expression of the AHR binding partner *ARNT* in either the parental or AHR-KO cells (**Figure 3D**).

### 3.5 AHR-KO clones show diminished basal CYP1A1 throughout pancreatic differentiation compared to parental cells

We next differentiated AHR-KO and parental hESCs into SC-islets using our 7-Stage pancreatic differentiation protocol (**Figure 4A**) to determine if AHR has a role in islet development. SC-islet cluster size was modestly increased in AHR-KO clone D8 on Day 31 compared to parental cells, but this effect was not seen in the other clones (**Figure 4B**). Qualitative examination of cluster morphology during differentiation showed no obvious differences between cells generated from any of the three AHR-KO clones versus the parental line (**Figure 4C**).

**Figure 4.**
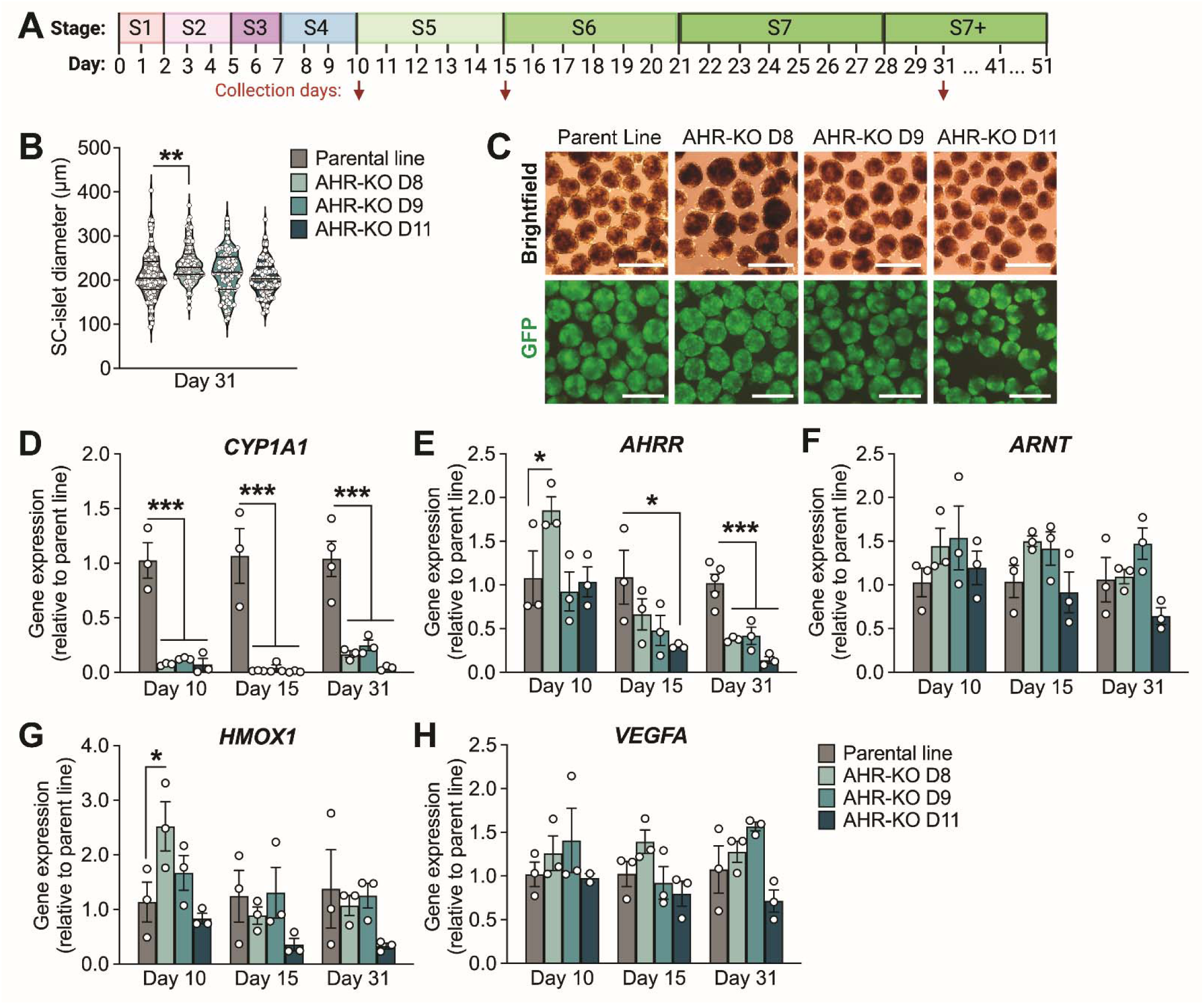
AHR-KO hESC clones generate morphologically normal SC-islets but with profoundly suppressed basal *CYP1A1* expression compared to parental cells. (A) INS-2A-EGFP parental hESCs and three AHR-KO clones (D8, D9, D11) were collected on Days 10 (Stage 4), 15 (Stage 5), and 31 (Stage 7+) of pancreatic differentiation for phenotyping. **(B)** SC-islet diameter was quantified on Day 31 of differentiation (n=1 biological replicate per cell line). (**C**) Representative brightfield and fluorescence images of INS-2A-EGFP parental cells and AHR-KO clones are shown on Day 31 of differentiation. Scale bars are 500 μm. **(D-H)** Expression of genes related to the AHR pathway and AHR-HIF1α crosstalk: (D) *CYP1A1,* (E) *AHRR*, (F) *ARNT*, (G) *HMOX1*, and (H) *VEGFA*. Gene expression is normalized to the parental cells at each timepoint. Data represent mean +/- SEM and individual data points represent biological replicates (n=3-4 per cell line). Asterisks indicate statistically significant differences relative to parental cells (*p<0.05, ***p<0.001), as determined by **(B)** one-way ANOVA with Dunnett post-hoc test, and **(D-H)** one-way ANOVA with Fisher LSD post-hoc test. **(A)** Schematic was made with BioRender.com.

Expression of AHR and HIF1α pathway target genes was measured at the end of Stage 4 (Day 10, pancreatic progenitor cells), end of Stage 5 (Day 15, pancreatic endocrine precursor cells) and mid-way through Stage 7+ (day 31, maturing pancreatic endocrine cells) (**Figure 4A**). Interestingly, basal *CYP1A1* expression was profoundly diminished in all AHR-KO clones compared to parental cells at all stages of differentiation assessed (mean of 11-fold, 46-fold, and 6.7-fold decrease across all clones on Days 10, 15, and 31 respectively; **Figure 4D**). *AHRR* expression was variable in the AHR-KO clones on Days 10 and 15 but was robustly decreased in all AHR-KO clones compared to parental cells on Day 31 (**Figure 4E**). There were no consistent differences between the AHR-KO clones and parental cells for levels of *ARNT, HMOX1,* or *VEGFA* on any of the days examined (**Figure 4F-H**).

### 3.6 SC-islets derived from AHR-KO clones show decreased G6PC2 expression and increased insulin secretion compared with parental cells at Stage 7

Key lineage markers of pancreatic endocrine cell differentiation were measured on Day 10 (*NGN3, PDX1),* Day 15 (*NGN3, PDX1, NKX6.1*) and Day 31 (*INS, GCG, G6PC2, MAFA, NKX6.1, SLC2A1*) (**Figure 4A**). *PDX1* was transiently reduced in all three AHR-KO clones compared to parental cells on Day 10 of differentiation but expression was restored by Day 15 (**Figure 5A,B**). There was no consistent effect of genotype on expression of *NGN3, NKX6.1, INS, GCG, SLC2A1,* or *MAFA* (**Figure 5A-C**). Notably, all AHR-KO clones showed a modest ∼2-fold decrease in *G6PC2* expression compared with parental cells on Day 31 (**Figure 5C**).

**Figure 5.**
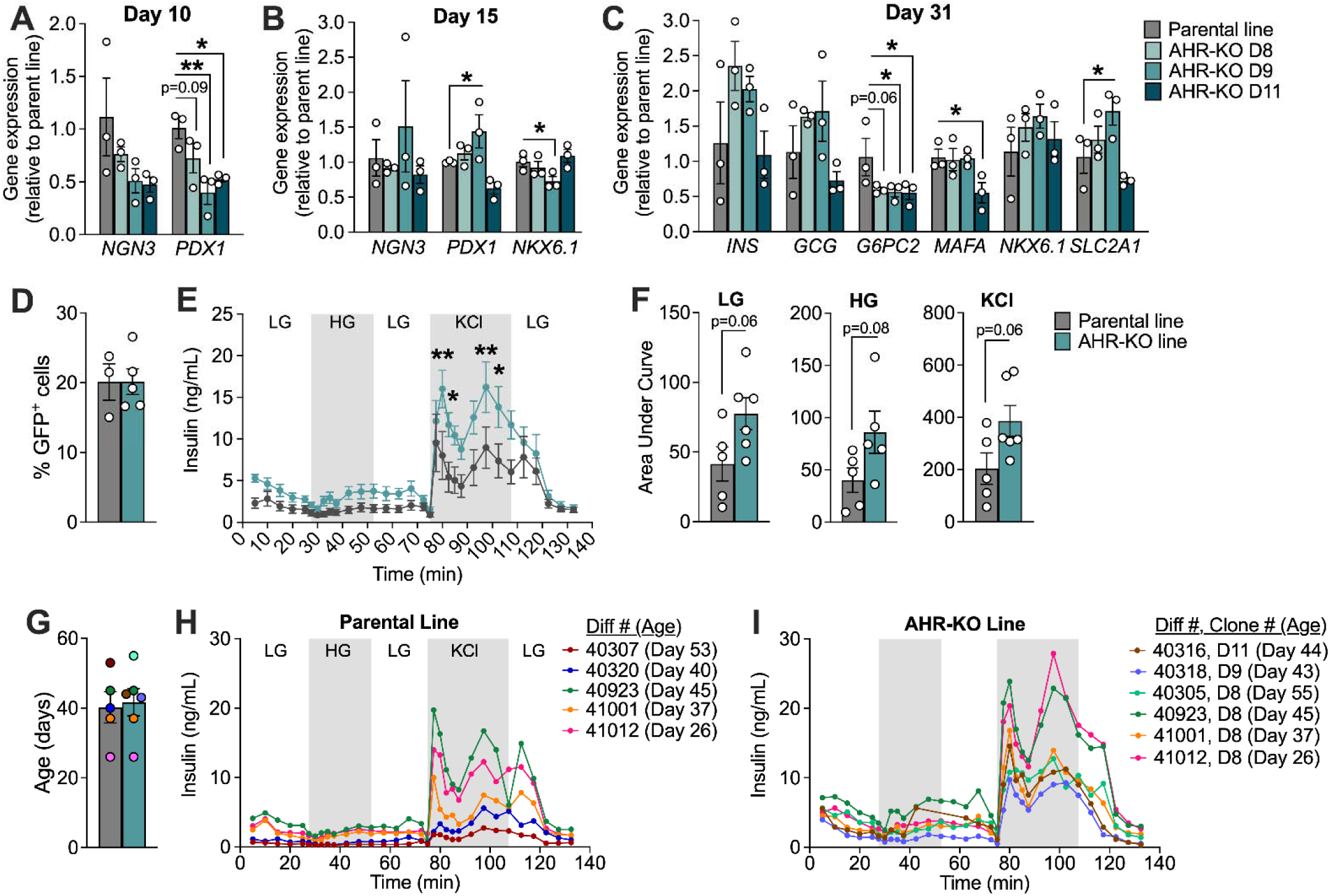
AHR-KO SC-islets have modestly decreased *G6PC2* expression and higher insulin secretion compared to parental cells. INS-2A-EGFP parental hESCs and AHR-KO clones D8, D9, and D11 were differentiated into SC-islets. See Figure 4A for schematic illustrating collection days. **(A-C)** Gene expression was measured for key markers of developing and mature pancreatic islets on **(A)** Day 10 (Stage 4), **(B)** Day 15 (Stage 5), and **(C)** Day 31 (Stage 7+) of differentiation (n=3 biological replicates per cell line at each timepoint). Gene expression is normalized to the parental cells at each timepoint. **(D)** Percentage of cells expressing GFP in SC-islets differentiated from the INS-2A-EGFP parental line (Day 40-52; n=3 biological replicates) and AHR-KO lines (Day 43-49; n=4 biological replicates). **(E,H,I)** Insulin secretion was measured in response to 2.5 mM low glucose (LG), 16.7 mM high glucose (HG), and 30 mM KCl during a dynamic perifusion analysis of SC-islets at Day 26-55 of differentiation (n=4-5 biological replicates for each genotype). **(F)** Area under the curve was calculated for the insulin response to LG, HG, and KCl. **(G)** Age (i.e., day of differentiation) of each biological replicate at the time of perifusion. **(E,F)** Perifusion data are plotted as mean +/- SEM and individual data points represent biological replicates. **(H,I)** Individual perifusion curves for each biological replicate are plotted separately for **(H)** the parental line and **(I)** AHR-KO line. Asterisks indicate statistically significant differences (*p<0.05, **p<0.01), as determined by **(A-C)** a one-way ANOVA with Fisher LSD post-hoc test, and **(E)** two-way repeated measures ANOVA with Sidak post-hoc test for perifusion curves (line graph), and **(F)** unpaired two-tailed t-test for AUCs (bar graphs).

There was no significant difference in the percentage of GFP^+^ cells between the INS-2A-EGFP parental line and AHR-KO clones on Day 40-52 of differentiation (**Figure 5D**). Yet, our dynamic perifusion assay showed that SC-islets generated from AHR-KO cells had significantly higher insulin secretion than the parent line at multiple time points during stimulation with 30 mM KCl (**Figure 5E**). Additionally, the overall area under the curve for insulin release under low glucose (2.8 mM), high glucose (16.7 mM), and KCl stimulation conditions trended towards being higher in AHR-KO cells compared to parental cells (p=0.06, p=0.08, p=0.06 respectively; **Figure 5E,F**). The different biological replicates used for perifusion analysis ranged in age from Day 26 to 55, but there was no difference in the average age of differentiation between genotypes (**Figure 5G**). Individual perifusion curves for each biological replicate are provided in **Figure 5H** (parental line) and **Figure 5I** (AHR-KO line).

## 4.0 DISCUSSION

There are thousands of diverse AHR ligands (e.g., dioxins, PAHs) polluting the environment^61^. Our study strongly suggests that exposure to exogenous AHR ligands during embryogenesis will activate AHR signalling in the developing human pancreas. We show that TCDD robustly activated AHR, as determined by induction of downstream targets *CYP1A1* and *AHRR*, as early as Day 5 of *in vitro* pancreatic differentiation―corresponding to induction of the primitive gut tube^62,63^―and consistently thereafter during the formation and maturation of pancreatic endocrine cells. Our data also point to a potential role for AHR in islet development; SC-islets generated from AHR-KO cells had modestly reduced *G6PC2* expression and higher insulin secretion compared to the parental line.

Our study focused mainly on the impact of acute AHR activation via 10 nM TCDD exposure for just 24 hours at each stage of pancreatic differentiation; our goal was not to maximally activate AHR, but to determine if AHR activation was possible at each stage of differentiation. Although there were no pronounced effects of acute TCDD exposure on key markers of pancreatic endocrine cell development, our data clearly demonstrate that the AHR pathway can be activated throughout human pancreas development. Additionally, we confirmed that continuous (72 hour) exposure to an AhR ligand at Stage 4 leads to sustained activation of AhR-target genes without altering *PDX1* or *NKX6.1* expression. Future studies should leverage this unique hESC model system to explore the impact of chronic AHR activation by exogenous ligands―both throughout the full differentiation and throughout individual stages―on pancreatic endocrine cell lineage commitment. Continuous AHR activation throughout the full differentiation would better replicate our mouse model in which we reported that chronic low-dose TCDD exposure throughout fetal development led to reduced β-cell mass in female pups at birth^48^.

We are also interested in understanding the basal role of AHR during pancreatic endocrine cell development (i.e., in the absence of a potent exogenous ligand like TCDD). Deletion of AHR in hESCs did not overtly impact the generation of GFP^+^ β-like cells *in vitro* or the expression of key hormones (*INS, GCG*) and β-cell identity genes (*NKX6.1*, *MAFA* and *SLC2A1*). However, we noted a modest but statistically significant decrease in gene expression of *G6PC2*—an enzyme primarily found in islets that opposes the action of glucokinase by converting glucose-6-phosphate into glucose—in Stage 7+ AHR-KO SC-islets. This coincided with a modest increase in glucose- and KCl-induced insulin secretion in SC-islets generated from AHR-KO cells. We did not normalize insulin secretion to total cell number or DNA content of the SC-islets, so it is possible that the increased insulin secretion is simply a reflection of higher cell number in SC-islets generated from AHR-KO versus parental cells. However, the improved insulin secretion in AHR-KO cells aligns with studies showing that downregulation of *G6PC2* may be a protective adaptation in islets^64–68^. For example, islet-specific *G6PC2* deletion in mice and in a human β-cell line also led to increased basal and stimulated insulin secretion^69,70^. *G6PC2* downregulation is also associated with reduced fasting glucose levels in rodents and humans^67,68,70–73^. Human islets from donors with T2D also have downregulated mRNA and protein expression of G6PC2^74,75^. Interestingly, TCDD treatment in Stage 7 modestly decreased *G6PC2* expression, which is consistent with previous data from our lab in SC-islets^26^; this also aligns with our mouse model in which female mice exposed to low-dose TCDD alongside high fat diet for 12 weeks *in vivo* had substantially reduced *G6pc2* expression in islets compared to vehicle-exposed mice fed either chow or high fat diet^76^. Taken together, these data point to a possible link between the AHR pathway and regulation of *G6PC2* expression. The role of the AHR in modulating *G6PC2* expression in islets should be further investigated.

Basal *CYP1A1* gene expression was significantly induced in parental hESCs starting at Day 15 of pancreatic differentiation (compared to Day 5) and was profoundly downregulated in all AHR-KO clones compared with the parental cell line at Days 10, 15, and 31. Unfortunately, we have been unable to find reliable CYP1A1 antibodies^25^ to determine if these changes in gene expression are also reflected at the protein level. Regardless, these dynamic changes in basal *CYP1A1* gene levels throughout differentiation in the parental, but not AHR-KO, cell line suggests that there are AHR ligands in the differentiation media. For example, the amino acid L-tryptophan and its metabolites are well-documented endogenous AHR ligands^77,78^; L-tryptophan has been shown to activate the AHR pathway in the human pancreatic cell line ECN90, human islets, and mouse islets^79^. However, we report significant *CYP1A1* induction starting at Day 15 (end of Stage 5), which does not coincide with changes in L-tryptophan concentration in the differentiation media; L-tryptophan increases from 0.025 mM in Stages 0-2 (RPMI 1640) to 0.078 mM in Stages 3-5 (DMEM HG). This suggests that there are other unidentified AHR ligands in the media, although this must be confirmed in future studies. Given that our stage-specific differentiation media composition is carefully designed to mimic the normal endogenous cues that pancreatic endocrine cells are exposed to throughout development, our findings point to a role of basal AHR activation in the developing endocrine pancreas. An additional explanation for basal upregulation of *CYP1A1* in the parental cell line is unliganded AHR activation. Studies have shown that unliganded AHR is capable of nucleocytoplasmic shuttling with potential roles in cell proliferation^80,81^. Further in-depth study of the AHR-KO line will hopefully provide insight into the complex biology of the AHR, both ligand-independent and ligand-activated, during differentiation. Additionally, our study focussed on canonical AHR target genes related to xenobiotic metabolism and AHR pathway autoregulation, but future studies should explore changes in gene expression with a more unbiased approach such as RNA-sequencing.

To our knowledge, we are the first to generate a human stem cell line with functional deletion of AHR. We thoroughly validated the AHR knock-out in three separate clones by confirming the absence of AHR protein, as well as the complete lack of *CYP1A1* and *AHRR* induction following exposure to the potent AHR ligand, TCDD. Given that hESCs can be differentiated into any cell type, the possible applications of this cell line are diverse. Importantly, AHR has diverse roles that extend beyond xenobiotic metabolism. There is extensive evidence that basal AHR activity contributes to development of diverse cell types and healthy maintenance of adult stem cell populations^17,22,24,82–91^. Additionally, AHR crosstalks with signaling pathways involved in controlling cell cycle, extracellular adhesion, lipid metabolism, circadian rhythm, inflammation and immune responses, and adaptation to hypoxia and oxidative stress^84,85,89,91–108^. Our novel AHR-KO hESC line thus allows for deeper exploration of AHR biology in diverse human cell lineages, such as the liver and gut. Future studies could also take advantage of the FKBP12^F36V^ degron tag to transiently deplete AHR, for example at a later stage of differentiation rather than from pluripotency. Application of the dTAG molecule induces rapid, reversible and selective degradation of FKBP12^F36V^ fusion proteins *in vitro* and *in vivo*^57^.

An interesting potential application of the AHR-KO hESC line is in islet transplantation research. The success of islet transplantation is currently hampered by compromised hypoxia signaling, which results in poor cell engraftment outcomes^109–111^. Since AHR and HIF1α compete for the same binding partner (ARNT), AHR activation may dampen hypoxia signaling^22,104,107^. We previously demonstrated that AHR-HIF1α crosstalk occurs in SC-islets and human donor islets; specifically, hypoxia exposure suppressed TCDD-mediated induction of *CYP1A1* expression^26^. We hypothesize that abolishing AHR competition for ARNT could enhance the hypoxia response post-transplant of SC-islets or other cell types and improve graft survival. Our current study shows that AHR deletion did not affect baseline expression of *VEGFA* at Days 10, 15, or 31 of differentiation *in vitro*, but future *in vivo* studies should test the ability of AHR-KO cells to mount a hypoxia response following transplantation.

In conclusion, our study suggests that AHR may be involved in regulating *G6PC2* and influencing insulin secretion in SC-islets. More detailed studies are needed to fully understand how both basal and ligand-activated AHR contribute to islet development and function. Our novel AHR-KO hESC line is a promising new tool to study the canonical and non-canonical roles of AHR in pancreatic islets and other cell lineages.

## AUTHOR CONTRIBUTIONS

NG, JEB, and FCL conceived the experimental design. FCL designed the AHR-KO construct. NG and FCL generated the AHR-KO construct. NG transfected and selected AHR-KO clones, performed TCDD experiments, and conducted genotyping and qPCR analysis. NG, FCL, EF, DZ, and BL differentiated hESCs. CN ran the Western blots. MPH conducted perifusion, flow cytometry, and *MAFA* and *G6pc2* qPCR analysis. BL conducted flow cytometry, the longer term TCDD/BaP study and TCDD dose response study, as well as qPCR analysis for several genes in Figure 1. NG, FCL, MPH, BL, and JEB analyzed the data. NG and JEB wrote the manuscript. All authors read and approved the manuscript.

## ACKNOWLEDGEMENTS

This research was supported by a Canadian Institutes of Health Research-Juvenile Diabetes Research Federation (CIHR-JDRF) Breakthrough Type 1 Diabetes (T1D) Team Grant (#ASD-173663/5-SRA-2020-1059-S-B) and Natural Sciences and Engineering Research Council (NSERC) Discovery Grant (#RGPIN-2017-06265). NG was supported by an NSERC Canadian Graduate Student-Doctoral (CGS-D) award and EF was supported by an NSERC-Collaborative Research and Training Experience (CREATE) award. JEB is supported by a Dorothy Killam Fellowship and an Early Researcher Award from the Ontario Government. We thank Drs. William Willmore and Jan Mennigan for thoughtful discussions over experimental design, Cameron Sinclair for qPCR primer design and validation, and Samantha Mar for support with sample collection.

The authors acknowledge that Carleton University is situated on the traditional, ancestral, and unceded territories of the Nehiyahwak, Nahikawe, Dakata, Lakota, Dene, and Metis Nations. The authors acknowledge that UBC and BC Children’s Hospital are situated on the traditional, ancestral, and unceded territories of the Coast Salish peoples—the S wxwú7mesh (Squamish), Səlı’lwəta /Selilwitulh (Tsleil-Waututh), and x məqk əyəm (Musqueam) Nations.

## DATA AVAILABILITY

Some or all datasets generated during and/or analyzed during the current study are not publicly available but are available from the corresponding author on reasonable request.

## Notes

https://data.mendeley.com/datasets/49jc6bkdt7/1

### Competing Interest Statement

The authors have declared no competing interest.

